# Heritability of de novo germline mutation reveals a contribution from paternal but not maternal genetic factors

**DOI:** 10.1101/2022.12.17.520885

**Authors:** Seongwon Hwang, Matthew D. C. Neville, Genomics England Research Consortium, Felix R. Day, Aylwyn Scally

## Abstract

De novo mutations (DNMs) in the germline have long been identified as a key element in the causes of developmental and other genetic disorders. Previous attempts to investigate genetic factors affecting DNMs have suffered from a lack of statistical power, due to the difficulty of obtaining a sufficient number of parent-offspring trios. Thus, the rare disease cohort of the UK’s 100k Genomes Project (100kGP), comprising more than 10,000 trios, represents an unprecedented opportunity to investigate the genetics of germline mutation. Here we estimate SNP heritability of DNM count in offspring, as a measure of the relative contribution of genetic factors to the variance of the trait, in a PCA-selected subset of the 100kGP cohort. We estimate separate SNP heritabilities for paternally and maternally transmitted mutations (based on parentally phased DNMs in offspring), computed using parental genetic variants at a range of minimum frequencies and a variety of methodologies. We estimate a heritability of 10-20% for paternal DNMs; by contrast, for maternal DNMs we find no significant evidence for non-zero heritability. We investigated the partitioning of heritability among genes with different expression profiles in different tissue or cell states, and found a relative heritability enrichment for genes expressed in gonadal tissues, particularly testis. Among germ cells in adult testes we observed relative enrichment of heritability in genes associated with the (undifferentiated) spermatogonial stem cell state.

## Introduction

Germline de novo mutations (DNMs) are DNA sequence mutations occurring within a generation on the lineage from parental to offspring zygote, and thus appearing as new genetic variants in all offspring cells. When they affect functional genomic regions they can cause adverse phenotypic effects (Veltman and Brunner, 2012) (Samocha et al., 2014), so understanding the factors contributing to germline mutation is important for the genetics of rare diseases and developmental disorders. However, DNMs are rare in the genome, and investigating them requires whole genome sequences from complete parent-offspring trios or larger pedigrees. Early studies of de novo point mutations in humans were therefore hindered by the cost and difficulty of whole genome sequencing, and the introduction of next-generation sequencing (NGS) technologies was crucial in overcoming these limitations (Koboldt et al., 2013). Over the last decade, the sizes of sequenced pedigree and trio cohorts have steadily increased, enabling DNM processes and the factors affecting them to be investigated with increasing statistical power.

Studies to date have focused on estimating the DNM rate in humans and the effects on it of parental age and genomic context (Kong et al., 2012) (Kessler et al., 2020) (Jónsson et al., 2017) (Goldmann et al., 2016). Cohorts with a typical parental age distribution reveal a mean DNM rate of 1.20×10^−8^ per nucleotide per generation, or about 70 DNMs per individual from both parents, with the majority coming from fathers. Older parents contribute more DNMs to offspring, with paternal age having roughly four times the effect of maternal age; for example in a study of 1,548 Icelanders (Jónsson et al., 2017) the number of DNMs from fathers increased by 1.51 per year, and from mothers only 0.37 per year. The causes of these age effects are somewhat unclear, since the prior model, in which the paternal effect was due to the greater number of cell divisions on the male germline (Crow, 2000) seems incompatible with observations such as that the male-female DNM ratio remains constant with age (Gao et al., 2019).

Other studies, both in trio cohorts and population genomic data, have detected the influence of local genomic factors such as histone modifications, recombination rate and replication timing on DNM rate (Francioli et al., 2015) (Carlson et al., 2018). DNMs are also found to cluster spatially on the genome, and factors determining the distribution, composition and age effect of clustered mutations appear to differ between sexes (Jónsson et al., 2017) (Goldmann et al., 2018).

Germline mutation processes have also been investigated by analysing the distribution of DNM rates by substitution type, referred to as the mutation spectrum, and its decomposition into distinct mutational signatures. A wide range of signatures have been observed in somatic mutations (Alexandrov et al., 2020), of which only two have been found to contribute substantially to germline mutation. Both have been associated with age-related mutagenesis (i.e. accumulating steadily over time) in somatic tissues, one seemingly driven by C>T mutations caused by spontaneous deamination of methylated cytosine (Rahbari et al., 2016). Evidence from population genetic datasets has also revealed that the base composition and wider spectrum of germline mutation have varied over time and between human populations (Narasimhan et al., 2017) (Harris and Pritchard, 2017). Again, the causes underlying these changes are unknown, and a variety of genetic and environmental factors have been proposed (Carlson, DeWitt and Harris, 2020).

Indeed, it remains an open question whether differences in germline mutation are driven primarily by environmental mutagens or endogenous genetic factors. Environmental factors might include the effects of diet, ionising radiation or reactive chemicals, temperature, or any mutagenic factors exogenous to the individual in which mutations arise. Effects on the DNM count and spectrum have been identified in fathers exposed to ionising radiation (Holtgrewe et al., 2018) or chemoreactive agents (Ton et al., 2018) (Kaplanis et al., 2022).

Genetic factors might include a population of mutator alleles, perhaps affecting the processes of DNA repair or replication, increasing or decreasing the likelihood of some or all categories of mutation throughout the germline or at some point in development or gametogenesis (Seplyarskiy et al., 2021). Previous studies have failed to find evidence either for extant mutator alleles of large effect in humans (Seoighe and Scally, 2017) or for the heritability of DNM count as a genetic trait (Jónsson et al., 2017), but their power was limited respectively by the removal of alleles with detectable effects through selection (Milligan, Amster and Sella, 2022) and small sample size. On a longer timescale, the mean germline mutation rate is known to differ between primate species (Chintalapati and Moorjani, 2020), and it has been argued that germline mutation rates across species are consistent with a balance between selection and drift (Lynch et al., 2016). These evolutionary considerations suggest that germline mutation in humans is or has been under genetic influence to a greater or lesser extent.

Here we investigate the heritability of germline DNM in a larger cohort than previously studied (13,949 parent-offspring trios), considering the number of DNMs passed by parents to offspring as a genetic trait. With a dataset of this size (903,525 DNMs, of which 241,063 were assigned to a parent of origin by (Kaplanis et al., 2022)) we are able to separately estimate the heritability of paternal and maternally derived mutations, and investigate the enrichment of heritability in genomic regions associated with tissue-specific or gametogenetic stage-specific gene expression.

Heritability is a quantitative measure of the genetic contribution to a trait in a specific study cohort. In particular, narrow-sense heritability is defined as the proportion of the observed trait variance accounted for by the additive effects of inherited genetic variants, and is often estimated using twin and family-based studies. However, a genome sequencing dataset such as this enables us to identify genotypes at specific loci and evaluate the contribution of observed genetic variation to heritability individually and in total – a quantity called SNP heritability. Since any estimate of heritability is contingent both on the cohort studied and on the methodology used to estimate it, our interest here is as much in the comparison between sexes as in its absolute value. We therefore consider the results of multiple commonly used estimation approaches, both those using the individual genotypes directly such as Genome-wide Complex Trait Analysis (GCTA) (Yang et al., 2011) or Linkage Disequilibrium Adjusted Kinships (LDAK) (Speed et al., 2012), and those based on SNP summary statistics such as LD score regression (LDSC) (Bulik-Sullivan et al., 2015). (The latter, although lower in statistical power, is often used in cases where SNP data are restricted or unavailable.)

## Results

We analysed whole-genome sequence data from the 100kGP rare disease cohort. In order to minimise bias due to population stratification when estimating heritability by GCTA (Krishna Kumar et al., 2016) we performed a principal components analysis (PCA) to identify population structure in the data, with the aim of selecting a more genetically homogeneous subset of the data. Based on this we restricted our subsequent analysis to a subset comprising a) individuals with self-identified ‘White British’ ethnicity; and, additionally, b) individuals with ‘Not Stated’ ethnicity lying in the same region of PCA space as those selected in (a). The resulting dataset comprised 7,985 individuals (Father) and 8,048 individuals (Mother) for the genotype.

DNMs were assigned to a parent of origin (phased) using nearby informative heterozygous sites as described in (Kaplanis et al., 2022). In each parental category (maternal and paternal) we took the number of phased DNMs assigned to that category in offspring as the phenotype of interest and estimated its SNP heritability using the corresponding parental genotypes as shown in Fig 1. We varied the threshold minor allele frequency (MAF) of potential explanatory SNPs from 0.0003 (rare variants) to 0.05 (common variants) to evaluate the effect of this on estimated heritability. Also, to account for age effects, in each parental category the parental age at conception was used as a covariate in the heritability analysis.

**Figure 1.**
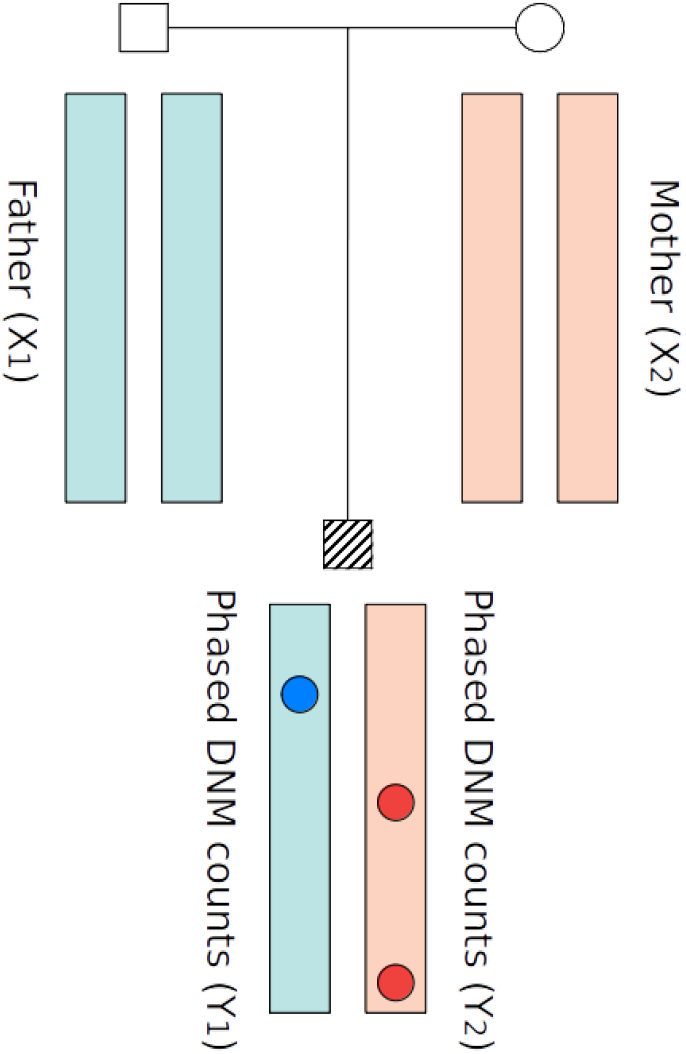
Diagram of family structure: blue and red circles represent DNMs in the child, assigned by phasing to a parental origin (maternal, red, or paternal, blue). The SNP heritability of paternal DNMs was estimated using corresponding paternal genotypes, and equivalently for maternal DNMs and maternal genotypes.

Estimates of SNP heritability were made using several methods, each involving different assumptions about the relationship between genotype and phenotype: GCTA, LDAK, HYBRID (an approach combining GCTA (Yang et al., 2011) and LDAK (Speed et al., 2012)), and LDSC (Bulik-Sullivan et al., 2015) (see Methods). The resulting estimates for both parental sexes are shown in Table 1, each for a range of allele frequency thresholds. Several overall comparisons and trends are apparent. In both parental categories, heritabilities estimated from summary statistics tend to be lower than those estimated from individual genotype (and we note that in principle the latter estimation method has more statistical power than the former).

**Table 1.**
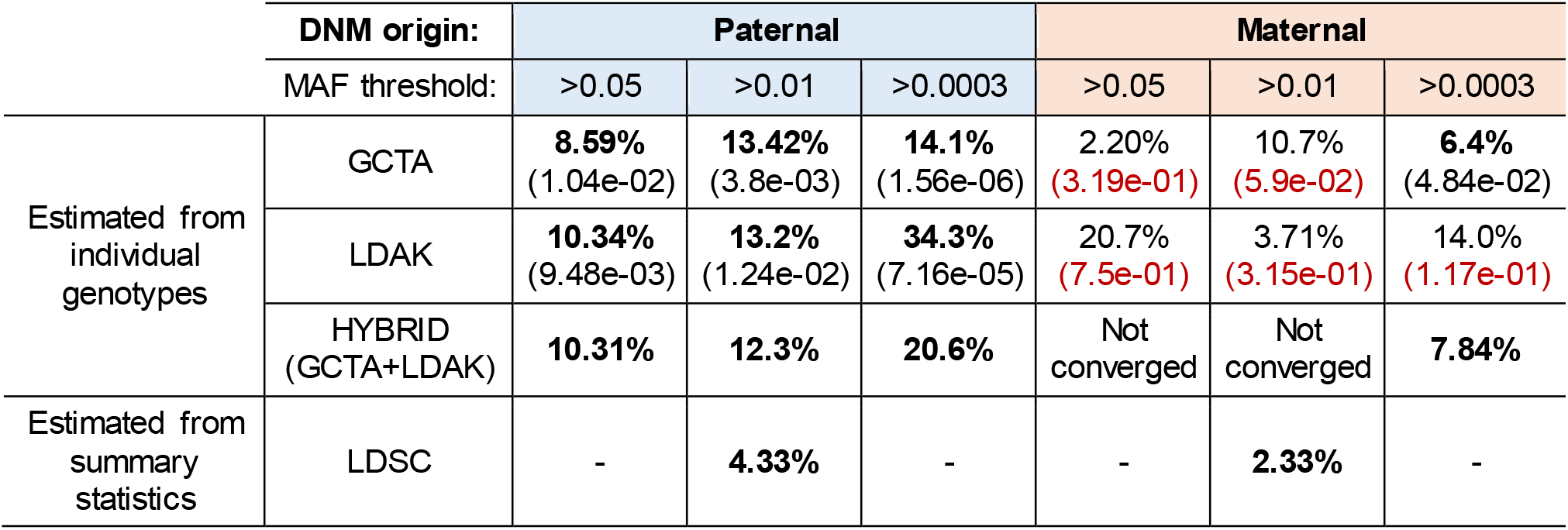
Heritability estimation of DNMs for both parents: Values in parentheses are p-values from the regression analysis for the random effect of variants; “Not converged” means restricted maximum likelihood estimates in the HYBRID model did not converge.

For paternal germline mutation, heritability estimates based on full genotype data range from around 10% to 20%, and increase with decreasing MAF threshold, from MAF 5% to 0.03%, by a factor of two or three depending on estimation method. This is consistent with the expectation that as more genetic variants are included, the proportion of phenotypic variance explained by them increases.

The picture is less clear for maternally phased DNMs, where most of the estimated heritability values are not significantly different from zero, and the HYBRID estimates for MAF thresholds 0.01 and 0.05 did not converge. The two exceptions, both at MAF threshold 0.03%, indicate a much lower heritability than the corresponding values for paternal mutation.

As shown in Table 2, we also estimated our statistical power to detect a non-zero heritability by the GCTA method. We estimated a power of more than 80% in case of paternal DNMs, but considerably less for maternal DNMs. This is likely due to the lower number of maternally phased DNMs in the analysis cohort (47,203 maternal compared to 163,715 paternal DNMs). Hence, we conclude that there is good evidence for a non-zero heritability of paternal de novo germline mutation in this cohort, with values ranging from 10-20% depending on estimation method and variants included. On the other hand, the impact of genetic influence on maternal DNM is far less clear.

**Table 2.** Statistical power calculation in GCTA model for both parents

| Statistical power |  | Father |  |  | Mother |  |  |
| --- | --- | --- | --- | --- | --- | --- | --- |
|  |  | >0.05 | >0.01 | >0.0003 | >0.05 | >0.01 | >0.0003 |
| Individual genotype | GCTA | **86.44%** | **92.46%** | **98.34%** | 12.00% | 70.93% | 48.27% |

In order to explore potential sources of observed heritability, we then investigated the contribution to heritability of variation in genes that are associated, via gene expression, with particular tissues (see Methods). The results of this analysis (Fig 2) suggest that heritability is particularly associated with variation in genes expressed in gonadal tissues (testis and vagina) compared to other tissues, but also nearby tissues such as cervix, prostate, bladder and suprapubic skin. In particular, SNP heritabilities in testis are enriched for both paternal and maternally derived DNMs, although we note that in general the number of maternal DNMs in a given category is substantially smaller, with correspondingly reduced power to discern enrichment.

**Figure 2.**
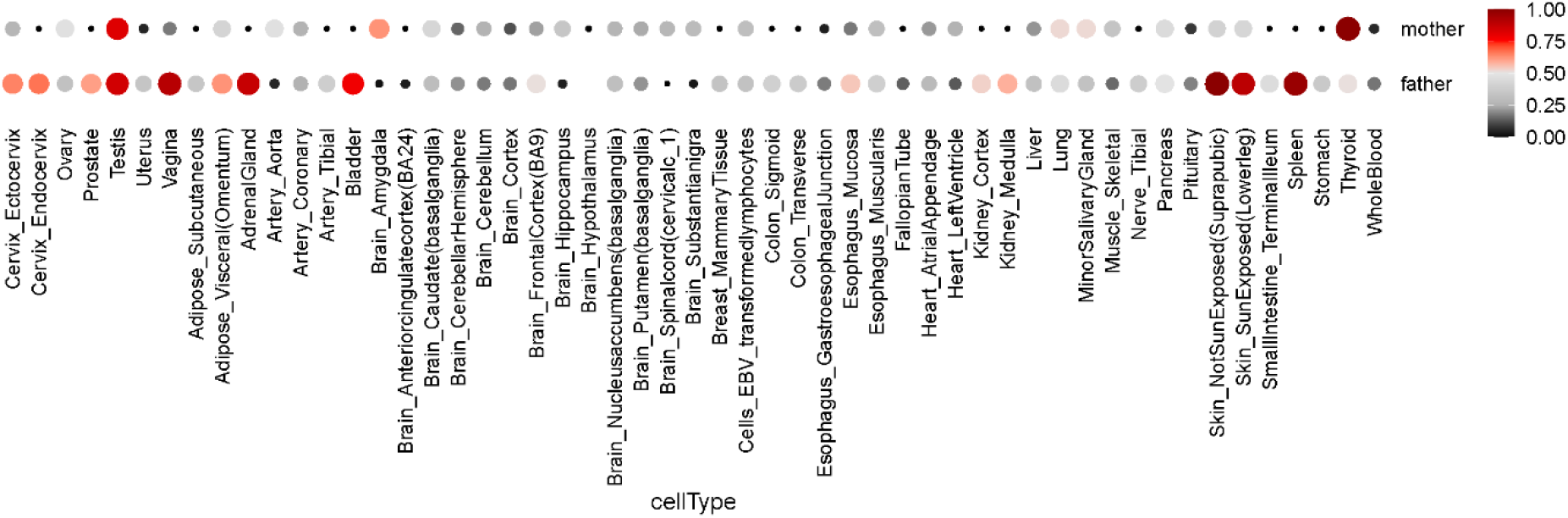
Heritability enrichment comparison among different tissues. The size of circle represents the relative heritability, and the degree of red colour also shows the relative heritability which is a mean per-SNP heritability relative to the max across all tissues.

To further investigate potential genetic factors mediated by biological processes in the germline, we performed a similar analysis partitioning heritability by gene clusters defined on the basis of expression in germline cells from adult testis samples (Guo et al., 2018). Our analysis revealed variable contributions to heritability from genes whose expression profiles were associated with different processes and cell states during spermatogenesis. The highest contribution came from variation in genes associated with undifferentiated spermatogonial stem cells (SSCs), compared with other clusters associated with processes such as differentiation, meiosis and spermatid formation. By contrast, one of the lowest contributions to heritability came from genes associated with meiosis, despite this being one of the larger clusters by number of genes.

As shown in Fig 3a, we see the same pattern regardless of the window size used to attribute SNPs to nearby genes, and also after normalising by the total number of SNPs attributed to each cluster (Fig 3b).

**Figure 3.**
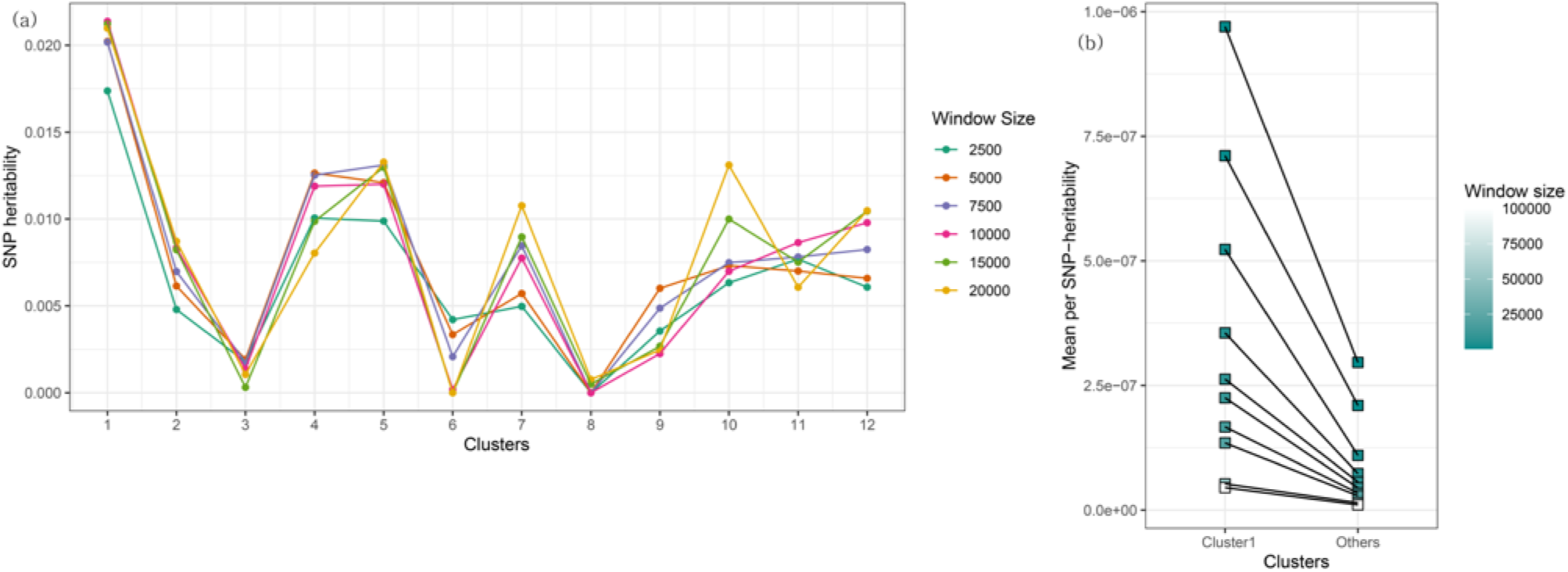
(a) Heritability enrichment comparison among seven biological processes during spermatogenesis. The 12 clusters used were associated with particular biological processes and stages of spermatogenesis by GO analysis in (Guo et al., 2018): 1: Transcription and signalling, 2: Differentiation and proliferation, 3: Meiosis; 4: Transcription and Signalling; 5: Mitochondrial translation; 6: Late pachynema; 7: Round spermatid & Elongating spermatid; 8, 9, 10: Elongating spermatid; 11, 12: Sperm. (b) Mean per SNP-heritability associated with the first cluster and with all other clusters during spermatogenesis: colour represents the window size used to attribute SNPs to nearby genes in a given cluster.

## Discussion

Genetic factors are known to play a role in somatic mutation, but prior to this study no significant influence had been detected on germline mutation (including an estimate of heritability in a much smaller cohort (Kessler et al., 2020)). The results presented here demonstrate that, for paternally inherited DNMs at least, genetic factors do affect germline mutation. Indeed, in the cohort studied they make a substantial contribution which is not dissimilar to that for other polygenic traits. However, the magnitude of any heritability estimate depends on the method used, including for example the number of SNPs used and the threshold on their frequency, as well as the genetic architecture of the trait and its interaction with environmental factors in the cohort studied. Of potentially greater interest here therefore is the marked difference in the estimated heritabilities of paternal and maternal DNM counts, which is consistent across multiple alternative methodologies. This presumably reflects physiological and developmental differences between the male and female germlines, and in fact our estimates are consistent with an absence of genetic influence on maternal germline mutation. However, given the lower rate of maternally transmitted DNMs, a larger cohort would be necessary in order to more precisely estimate a heritability for this trait. Similarly, discerning potential genetic influences on other mutational phenotypes such as spectrum, age effect or clustering will be possible in larger study cohorts drawn from this and other populations.

The fact that estimated heritability is, as might be expected, relatively enriched in genes associated with gonadal tissues and other tissues proximal to the germline, provides confidence in the detected signal of heritability itself (in the sense that the observed variation is genuinely associated with germline mutation). It also encourages us that partitioning heritability in this cohort by associated tissue or cell type is feasible and potentially informative, for paternal DNMs at least. Our finding that within the male germline there is enrichment in genes associated with the SSC state is consistent with the fact that in males the germline spends the majority of its existence in this state. In particular, based on observed markers (Guo et al., 2018), the SSC category in our analysis includes both active and seemingly quiescent (non-dividing) germline cells. Thus, the genetic factors we identify could be associated with either replicative or non-replicative mutation processes, or a combination of both. It is also notable that a low degree of relative enrichment comes from variation in meiosis-associated genes. Further investigation, again perhaps in larger cohorts, may discern individual genetic factors such as particular genes or pathways.

One issue to consider is whether any bias comes from the fact that offspring in the studied cohort are affected by rare genetic disease. For example, a higher mean DNM rate in affected cases than in unaffected cases could plausibly arise through correlation with paternal age or the ascertainment constraint of having a DNM with functional consequences (without any direct or causative connection with mutation processes). Therefore, it is conceivable a priori that details of our findings may have been affected. However, the cohort studied here covers over 190 different rare diseases. If most of these are due to causative mutations at specific developmental or other functional loci, we can reasonably expect the cohort to otherwise reflect the wider population in terms of genomic and mutational phenomena, including those investigated here. Therefore, we do not expect our estimates of heritability and its variation between sexes to be substantially biased by this aspect of the data.

Much remains unclear about the biological processes affecting germline mutation in humans and other organisms, and the causes of mutation rate change over evolutionary timescales. Recent studies have estimated similar values across multiple mammalian species for certain mutational parameters, such as the paternal age effect and the male/female ratio of mutation rates (de Manuel, Wu and Przeworski, 2022). Given the variety of environments involved, it may be that these similarities are due to shared genetic or other endogenous factors in mammalian biology, and investigation in a non-human primate cohort, if feasible, would be informative about this. Furthermore, and perhaps more speculatively, the characterisation of genetic factors affecting germline mutation today in humans or other primates raises the possibility of estimating the genetic contribution to mutation rates in ancestral species and populations. Notwithstanding the important (and perhaps unmeasurable) influence of ancestral environments, and the many issues in applying polygenetic scores as cross-cohort predictors of trait values, this might be a way to improve branch length and speciation time estimates within the great apes and other primate groups.

## Materials and methods

### Data

Analyses were carried out on a dataset of whole-genome sequences from 13,949 parent-offspring trio samples collected as part of the 100,000 Genomes Project Rare Disease cohort. All sequencing was carried by one sequence provider, using the Illumina platform, to a mean depth of 32x. Thus, little batch effect in 100kGP is expected to exist in our dataset. We started from a dataset of 903,525 de novo single-nucleotide variant calls (dnSNVs) (detailed in (Kaplanis et al., 2022)). After filtering to remove 20 hyper-mutated and other outlier individuals by looking at the z-score of DNM counts, this resulted in a dataset comprising 210,918 dnSNVs called in 10,749 trios, which were then phased by parent of origin (using reads and nearby informative heterozygous sites). Insertions and deletions (indels) were excluded from subsequent analyses.

### Heritability calculation

These variants were then used to compute the SNP heritability of offspring DNM count. We standardise the phenotype (phased DNM count) by computing a z-score for each proband, subtracting the mean phased DNM count across probands and then dividing these zero-centred values by the standard deviation of phased DNM count.

Some further filtering of genomic variants was applied before inclusion in the heritability calculation. We only included variants that have “PASS” in filters within VCFv4.2 format such as ‘q10’ if quality at this site is below 10 and ‘s50’ if less than 50% of samples have data. If the site passes all filters, “PASS”. Furthermore, in certain analyses we varied the minimum MAF, to select either common variants only (MAF 0.05 or 0.01) or both rare (MAF 0.0003) and common variants. The lowest (rare) MAF threshold was equivalent to the criterion that about 5 people among 8,000 participants have the variant. When more than 10% among all participants had a missing genotype at a certain variant, this variant was filtered out. In addition, we filtered out multi-allelic variants so that we only have a maximum of 2 alleles at a variant.

We estimated SNP heritability using several tools, each of which has different modelling assumptions about the random effect sizes of genotypes on the phenotype. Firstly, we used GCTA, which assumes the random contribution of each SNP to the phenotype in GCTA is identical across all SNPs. GCTA requires that all SNPs used be in linkage equilibrium, so we pruned our SNPs (using PLINK variant pruning) according to the criterion that pairs of variants in a window with r^2^ correlation greater than the threshold are greedily pruned from the window until no such pairs remain. We set up that the window size is 50 kilobases, step size is 10 variant count, and r^2^ threshold is 0.5. Furthermore, GCTA expects no genetic stratification in the population, otherwise maximum likelihood estimates computed by GCTA are biased (Krishna Kumar et al., 2016). Thus, we mitigated this bias by controlling for genetic population structure in the sampled data as represented in a PCA analysis, as described in the Results. In carrying out this PCA we filtered variants based on coverage, MAF, Linkage Disequilibrium (LD) pruning and other criteria.

We also estimated SNP heritability using LDAK, which assumes the random effect size for SNP is dependent on its allele frequency and the LDAK weightings of the SNP (Zhang et al., 2021). When SNPs are in regions of high LD, their LDAK weightings tend to be lower and the weightings are calculated by adding our genotype data into LDAK. Since LDAK takes LD structure into account by itself, there is no need to prune our dataset before running it. Although both the GCTA and LDAK methods use the individual genotype data for estimation of heritability, the LDAK authors recommend using a hybrid method combining GCTA and LDAK if there is a big difference between the estimates from these two methods.

For additional comparison with estimates for other traits, we also estimated SNP heritability using LDSC, a method based on summary-statistics. To generate suitable summary statistics in our dataset, we first performed GWAS for each trait (parental and maternal phased DNM counts) using a linear regression model with parental age as a covariate.

### Heritability enrichment

Besides the estimation of total SNP heritability with varying MAF thresholds, we also investigated the enrichment of heritability in specific tissues based on gene expression. We obtained genome-wide expression data for multiple tissues from GTEx (‘The Genotype-Tissue Expression (GTEx) project.’, 2013), and compiled a list of tissue-specific genes for each tissue, by computing t-statistics comparing the expression level of genes in a tissue of interest with that in other tissues, using the same approach as (Finucane et al., 2018). Genes in the top 10% by t-statistic for expression associated with a given tissue were considered to be associated with that tissue. For each tissue we then compiled a set of associated SNPs by including all SNPs within a certain distance of genes associated with that tissue (Fig 4). Noting that most of the GWAS-indicated SNPs were located in non-coding regions, we defined windows based on the transcription start site (TSS) rather than exonic region of each gene, and used a 100-kb window size, which was found to yield robust signals of heritability enrichment in (Finucane et al., 2018).

**Figure 4.**
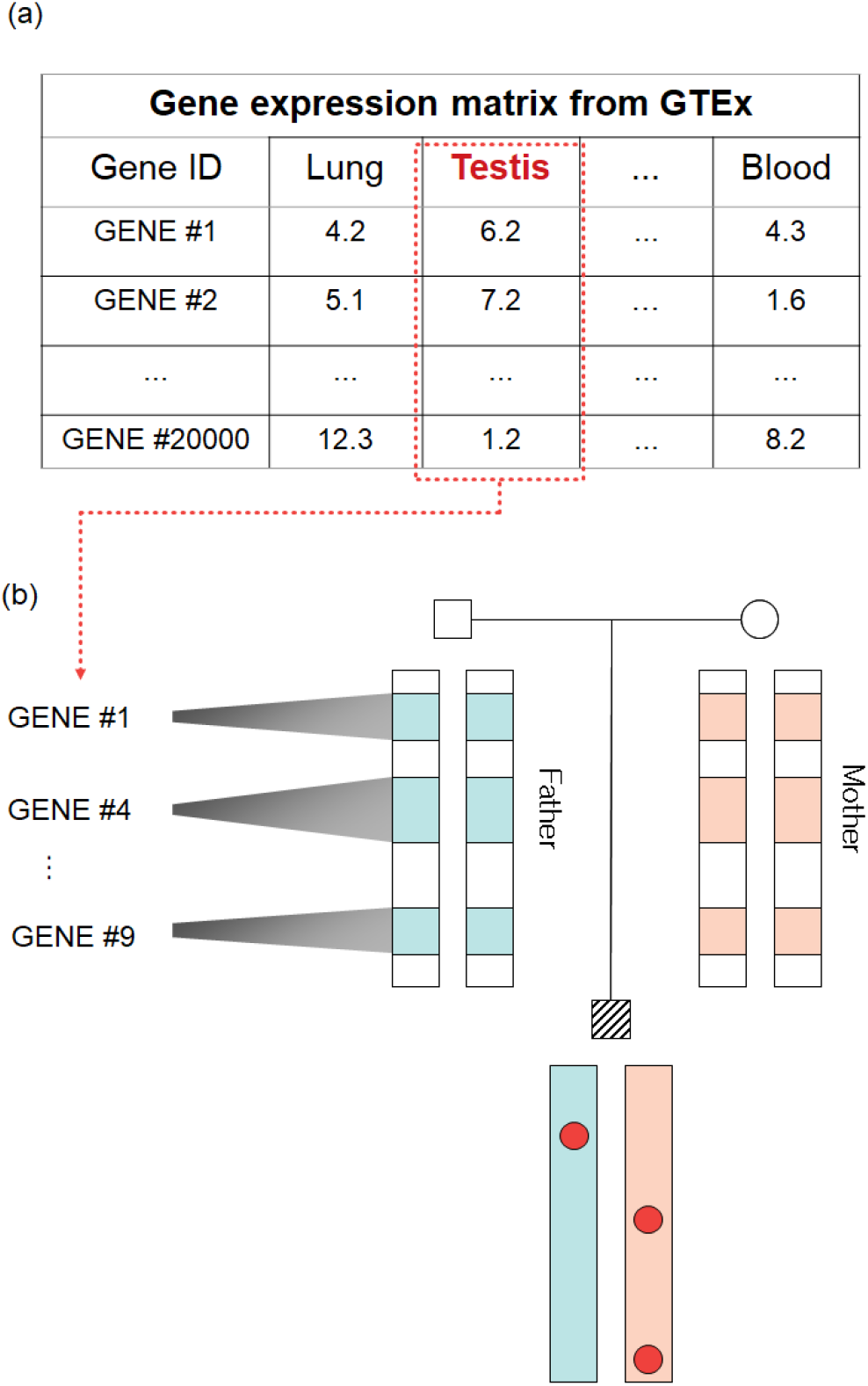
Schematic of tissue-specific heritability enrichment analysis: (a) Identification of tissue-specific genes based on expression profiles. (b) Selection of associated SNPs, near tissue-specific genes, used to estimate heritability paternal and maternal phased DNM counts.

Then for each tissue we estimated the mean per SNP-heritability of DNM count using only the relevant tissue-associated SNPs, in order to compare their relative contributions to heritability (Fig 4).

We carried out a similar analysis to identify genomic regions associated with biological processes during spermatogenesis. In this case we obtained gene expression data for spermatogonial cells which were classified into seven different clusters representing transcription/developmental states (or groups of states) by (Guo et al., 2018). T-statistics for the association of genes with each cluster were then inferred as above, and a heritability enrichment analysis carried out on nearby SNPs. To further investigate if genes associated with cluster 1, representing the SSC stage of spermatogenesis, make a particular contribution to the heritability, we also compared mean per SNP-heritability between this cluster and another category comprising genes associated to all other clusters.

## Acknowledgements

We thank Raheleh Rahbari for useful discussions. This research was made possible through access to the data and findings generated by the 100,000 Genomes Project. The 100,000 Genomes Project is managed by Genomics England Limited (a wholly owned company of the Department of Health and Social Care). The 100,000 Genomes Project is funded by the National Institute for Health Research and NHS England. The Wellcome Trust, Cancer Research UK and the Medical Research Council (MC_UU_00006/2) have also funded research infrastructure. The 100,000 Genomes Project uses data provided by patients and collected by the National Health Service as part of their care and support.

## Supplementary figures

**Supplementary Figure 1.**
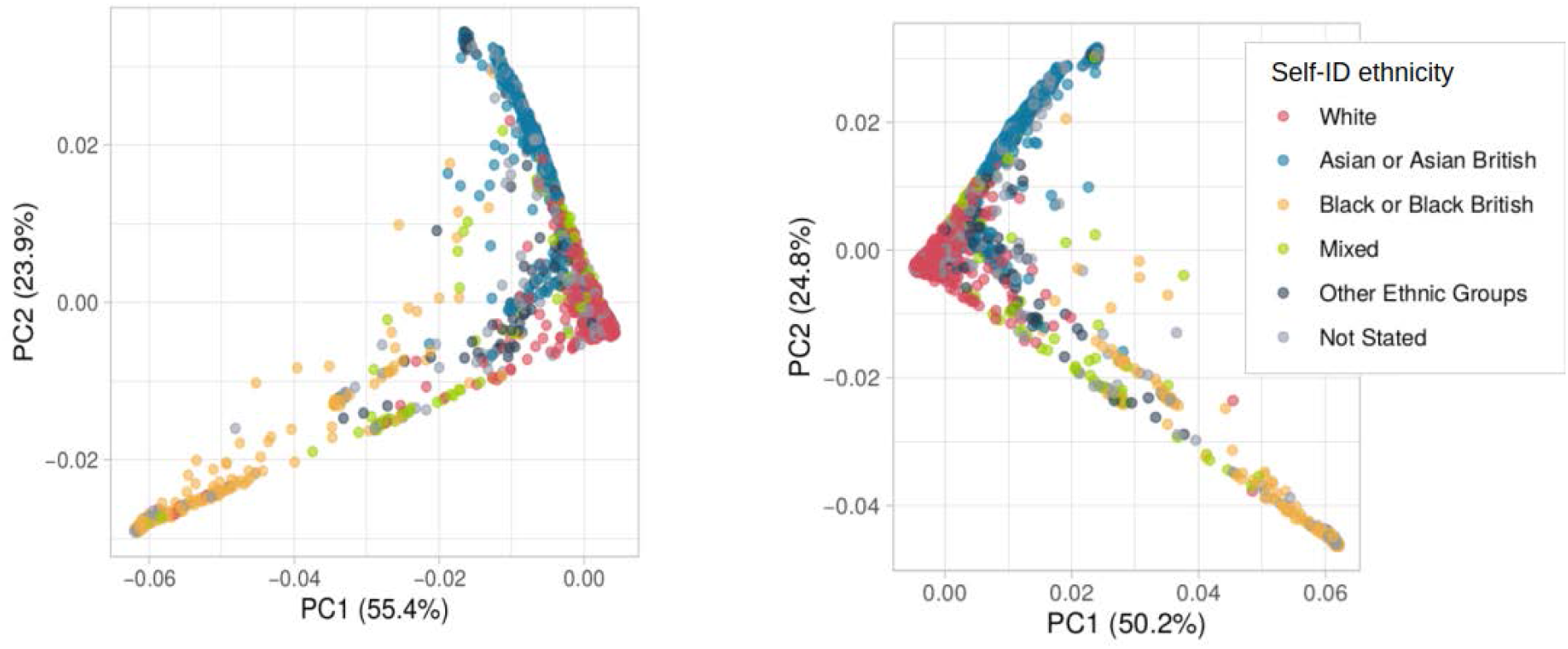
PCA plot for genotypes of each parent: the left is for father and the right is for mother. Colours were assigned by the race information from all subjects

**Supplementary Figure 2.**
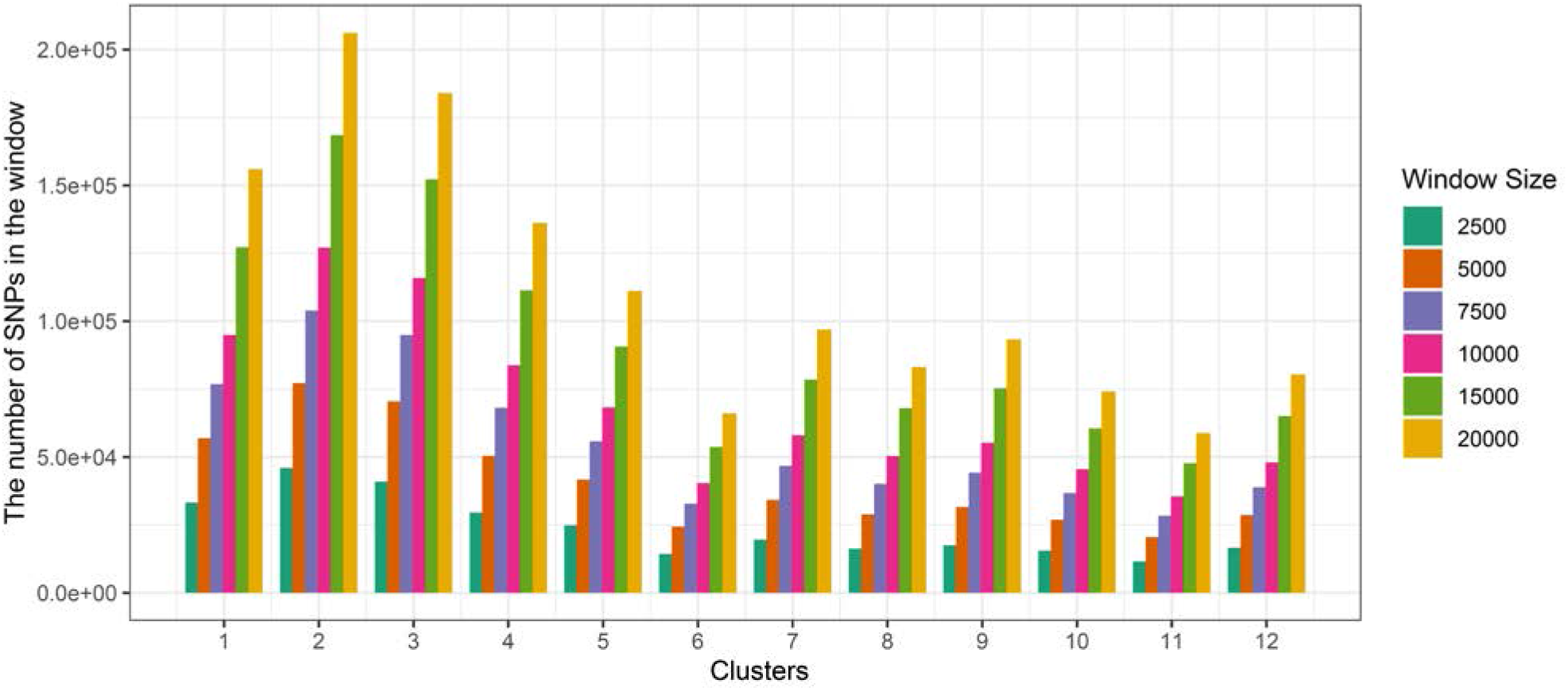
The number of SNPs within the window at each cluster

## References

Alexandrov, L. B. et al. (2020) ‘The repertoire of mutational signatures in human cancer’, Nature. Nature Publishing Group, 578(7793), pp. 94–101.

Bulik-Sullivan, B. K. et al. (2015) ‘LD Score regression distinguishes confounding from polygenicity in genome-wide association studies’, Nature Genetics, 47(3), pp. 291–295. doi: 10.1038/ng.3211.

Carlson, J. et al. (2018) ‘Extremely rare variants reveal patterns of germline mutation rate heterogeneity in humans’, Nature communications. Nature Publishing Group, 9(1), pp. 1–13.

Carlson, J., DeWitt, W. S. and Harris, K. (2020) ‘Inferring evolutionary dynamics of mutation rates through the lens of mutation spectrum variation’, Current Opinion in Genetics and Development. Elsevier Ltd, 62, pp. 50–57. doi: 10.1016/j.gde.2020.05.024.

Chintalapati, M. and Moorjani, P. (2020) ‘Evolution of the mutation rate across primates’, Current opinion in genetics & development. Elsevier, 62, pp. 58–64.

Crow, J. F. (2000) ‘The origins, patterns and implications of human spontaneous mutation’, Nature Reviews Genetics. Nature Publishing Group, 1(1), pp. 40–47.

Finucane, H. K. et al. (2018) ‘Heritability enrichment of specifically expressed genes identifies disease-relevant tissues and cell types’, Nature genetics. Nature Publishing Group, 50(4), pp. 621– 629.

Francioli, L. C. et al. (2015) ‘Genome-wide patterns and properties of de novo mutations in humans’, Nature Genetics. doi: 10.1038/ng.3292.

Gao, Z. et al. (2019) ‘Overlooked roles of DNA damage and maternal age in generating human germline mutations’, Proceedings of the National Academy of Sciences. National Acad Sciences, 116(19), pp. 9491–9500.

Goldmann, J. M. et al. (2016) ‘Parent-of-origin-specific signatures of de novo mutations’, Nature genetics. Nature Publishing Group, 48(8), pp. 935–939.

Goldmann, J. M. et al. (2018) ‘Germline de novo mutation clusters arise during oocyte aging in genomic regions with high double-strand-break incidence’, Nature genetics. Nature Publishing Group, 50(4), pp. 487–492.

Guo, J. et al. (2018) ‘The adult human testis transcriptional cell atlas’, Cell research. Nature Publishing Group, 28(12), pp. 1141–1157.

Harris, K. and Pritchard, J. K. (2017) ‘Rapid evolution of the human mutation spectrum’, Elife. eLife Sciences Publications Limited, 6, p. e24284.

Holtgrewe, M. et al. (2018) ‘Multisite de novo mutations in human offspring after paternal exposure to ionizing radiation’, Scientific reports. Nature Publishing Group, 8(1), pp. 1–5.

Jónsson, H. et al. (2017) ‘Parental influence on human germline de novo mutations in 1,548 trios from Iceland’, Nature. doi: 10.1038/nature24018.

Kaplanis, J. et al. (2022) ‘Genetic and chemotherapeutic influences on germline hypermutation’, Nature, 605(7910), pp. 503–508. doi: 10.1038/s41586-022-04712-2.

Kessler, M. D. et al. (2020) ‘De novo mutations across 1,465 diverse genomes reveal mutational insights and reductions in the Amish founder population’, Proceedings of the National Academy of Sciences of the United States of America. doi: 10.1073/pnas.1902766117.

Koboldt, D. C. et al. (2013) ‘The next-generation sequencing revolution and its impact on genomics’, Cell. Elsevier, 155(1), pp. 27–38.

Kong, A. et al. (2012) ‘Rate of de novo mutations and the importance of father’s age to disease risk’, Nature. Nature Publishing Group, 488(7412), pp. 471–475.

Krishna Kumar, S. et al. (2016) ‘Limitations of GCTA as a solution to the missing heritability problem’, Proceedings of the National Academy of Sciences. National Acad Sciences, 113(1), pp. E61–E70.

Lynch, M. et al. (2016) ‘Genetic drift, selection and the evolution of the mutation rate’, Nature Reviews Genetics. Nature Publishing Group, 17(11), pp. 704–714.

de Manuel, M., Wu, F. L. and Przeworski, M. (2022) ‘A paternal bias in germline mutation is widespread across amniotes and can arise independently of cell divisions’, bioRxiv. Cold Spring Harbor Laboratory.

Milligan, W. R., Amster, G. and Sella, G. (no date) ‘The impact of genetic modifiers on variation in germline mutation rates within and among human populations’. doi: 10.1101/2021.08.25.457718.

Narasimhan, V. M. et al. (2017) ‘Estimating the human mutation rate from autozygous segments reveals population differences in human mutational processes’, Nature communications. Nature Publishing Group, 8(1), pp. 1–7.

Rahbari, R. et al. (2016) ‘Timing, rates and spectra of human germline mutation’, Nature genetics. Nature Publishing Group, 48(2), pp. 126–133.

Samocha, K. E. et al. (2014) ‘A framework for the interpretation of de novo mutation in human disease’, Nature genetics. Nature Publishing Group, 46(9), pp. 944–950.

Seoighe, C. and Scally, A. (2017) ‘Inference of candidate germline mutator loci in humans from genome-wide haplotype data’, PLoS genetics. Public Library of Science San Francisco, CA USA, 13(1), p. e1006549.

Seplyarskiy, V. B. et al. (2021) ‘Population sequencing data reveal a compendium of mutational processes in the human germ line’, Science. American Association for the Advancement of Science, 373(6558), pp. 1030–1035.

Speed, D. et al. (2012) ‘Improved heritability estimation from genome-wide SNPs’, The American Journal of Human Genetics. Elsevier, 91(6), pp. 1011–1021.

‘The Genotype-Tissue Expression (GTEx) project.’ (2013) Nature genetics. United States, 45(6), pp. 580–585. doi: 10.1038/ng.2653.

Ton, N. D. et al. (2018) ‘Whole genome sequencing and mutation rate analysis of trios with paternal dioxin exposure’, Human Mutation. Wiley Online Library, 39(10), pp. 1384–1392.

Veltman, J. A. and Brunner, H. G. (2012) ‘De novo mutations in human genetic disease’, Nature Reviews Genetics. doi: 10.1038/nrg3241.

Yang, J. et al. (2011) ‘GCTA: a tool for genome-wide complex trait analysis.’, American journal of human genetics. United States, 88(1), pp. 76–82. doi: 10.1016/j.ajhg.2010.11.011.

Zhang, Q. et al. (2021) ‘Improved genetic prediction of complex traits from individual-level data or summary statistics’, Nature communications. Nature Publishing Group, 12(1), pp. 1–9.

